# Regulatory divergence and functional diversification of a c-di-GMP-controlled sigma factor in Actinomycetota

**DOI:** 10.64898/2026.01.14.699351

**Authors:** Joee D. Denis, Govind Chandra, Justine J. Choi, Yves V. Brun, Kelley A. Gallagher

## Abstract

Members of the σ^28^ family of alternative σ factors typically regulate genes involved in flagellar biosynthesis. However, the only member of the σ^28^ family present in the actinobacterial genus *Streptomyces*, WhiG, controls the differentiation of aerial hyphae into spores. WhiG activity is regulated by the second messenger c-di-GMP, which arms its cognate anti-σ, RsiG, to bind and sequester the σ factor. Understanding WhiG evolution can thus shed light on the diversity of processes regulated by c-di-GMP across Phylum Actinomycetota. Members of Actinomycetota comprise highly diverse filamentous and unicellular, flagellated and non-flagellated species, and the actinobacterial ancestor is predicted to have been motile. Here, our systematic analysis reveals that WhiG homologs are broadly distributed throughout the Actinomycetota and form two distinct clades: WhiG1, whose members have retained the ancestral association with the flagellar biosynthesis cluster and are not regulated by an RsiG anti-σ, and WhiG2, whose members are typically regulated by RsiG via c-di-GMP. Bioinformatic analysis of their target regulons suggests that this σ factor has significantly diversified in function to control various processes, including motility, chemotaxis, Type IV pili synthesis, sporulation, antibiotic biosynthesis, and c-di-GMP metabolism across diverse actinobacterial species. These findings highlight a phylogenetic split in the regulation and function of this key transcription factor throughout the phylum.

**Importance:** Mounting global responses to dynamic environmental conditions is a crucial function that bacterial cells must perform. Global regulatory networks are most often studied in individual species, however, understanding how regulatory networks have evolved in distinct bacterial lineages remains an outstanding question. The alternative σ factor WhiG, which is found in the actinobacterial genus *Streptomyces*, is a dedicated sporulation σ, yet has long been known to have a close evolutionary relationship to σ factors that are responsible for regulation of flagellar biosynthesis. Analysis of the distribution of WhiG in Phylum Actinomycetota reveals that homologs typically co-occur with either a flagellar cluster or the c-di-GMP-binding anti-σ RsiG. Functional predictions of WhiG target genes across the Actinomycetota further reveal that diverse biological processes are controlled by these transcription factors, most notably motility, chemotaxis, Type IV pili biosynthesis, regulation of gene expression, c-di-GMP metabolism, and specialized metabolite biosynthesis.

## Introduction

The nucleotide second messenger 3’, 5’-cyclic diguanylic acid (c-di-GMP) plays a critical role in coordinating global cellular responses to environmental changes (1). Cellular c-di-GMP levels are tightly regulated by the action of diguanylate cyclases (DGCs), which synthesize c-di-GMP, and phosphodiesterases (PDEs), which degrade it (2–4). DGCs and PDEs often contain sensory domains that respond to specific stimuli, modulating c-di-GMP levels through changes in their activity (5). Cellular levels of c-di-GMP are detected by effector proteins, which upon binding to c-di-GMP, can modulate the activity of downstream target systems involved in fundamental microbial behaviors such as motility, virulence, and biofilm formation (1). Homologs of DGCs and PDEs can be found in all major bacterial phyla (6). Despite this near ubiquity across the bacterial domain, most research has focused on Gram-negative bacteria, particularly within the Phylum Pseudomonadota, which has limited our understanding of the role of c-di-GMP in bacterial physiology from diverse taxa. In the filamentous bacterial genus *Streptomyces*, which resides in the phylogenetically distant Phylum Actinomycetota, c-di-GMP governs complex morphological development, including the formation of vegetative mycelium followed by aerial mycelium production and sporulation (7–9).

We previously showed that c-di-GMP regulates *Streptomyces* development by controlling activity of the σ factor WhiG (10). WhiG is a sporulation-specific σ factor that regulates expression of three known target genes in the model species *Streptomyces venezuelae*: the developmental regulators *whiH* (*vnz_27205*) and *whiI* (*vnz_28820*), and a poorly-conserved membrane protein of unknown function (*vnz_15005*) (10). *whiH* and *whiI* are also confirmed WhiG targets in the species *Streptomyces coelicolor* (11, 12). WhiH is a GntR-family transcriptional regulator (11), and WhiI is an orphan response regulator that forms a functional heterodimer with a second orphan response regulator called BldM (13). Both WhiH and WhiI are expressed during the late stages of *Streptomyces* development (10, 13, 14) and are essential for the proper maturation of aerial hyphae into spores. The precise timing of WhiG activity is thus crucial for proper development. The activity of WhiG is controlled post-translationally by a cognate anti-σ factor, RsiG. To interact with WhiG, RsiG must bind a partially intercalated dimer of c-di-GMP via two E(X)_3_S(X)_2_R(X)_3_Q(X)_3_D repeat motifs, one on each helix of an antiparallel coiled-coil. When bound to RsiG, WhiG is sequestered, leaving its set of target genes off. Cellular levels of c-di-GMP in *S. venezuelae* are high during vegetative growth and drop prior to sporulation (15). Therefore, the relatively high cellular levels of c-di-GMP during vegetative growth favors RsiG-(c-di-GMP)-WhiG complex formation (Fig 1). Later in the life cycle, c-di-GMP levels drop, the complex disassociates and WhiG can then direct RNA polymerase (RNAP) to transcribe its set of target genes (10, 16).

**Fig 1.**
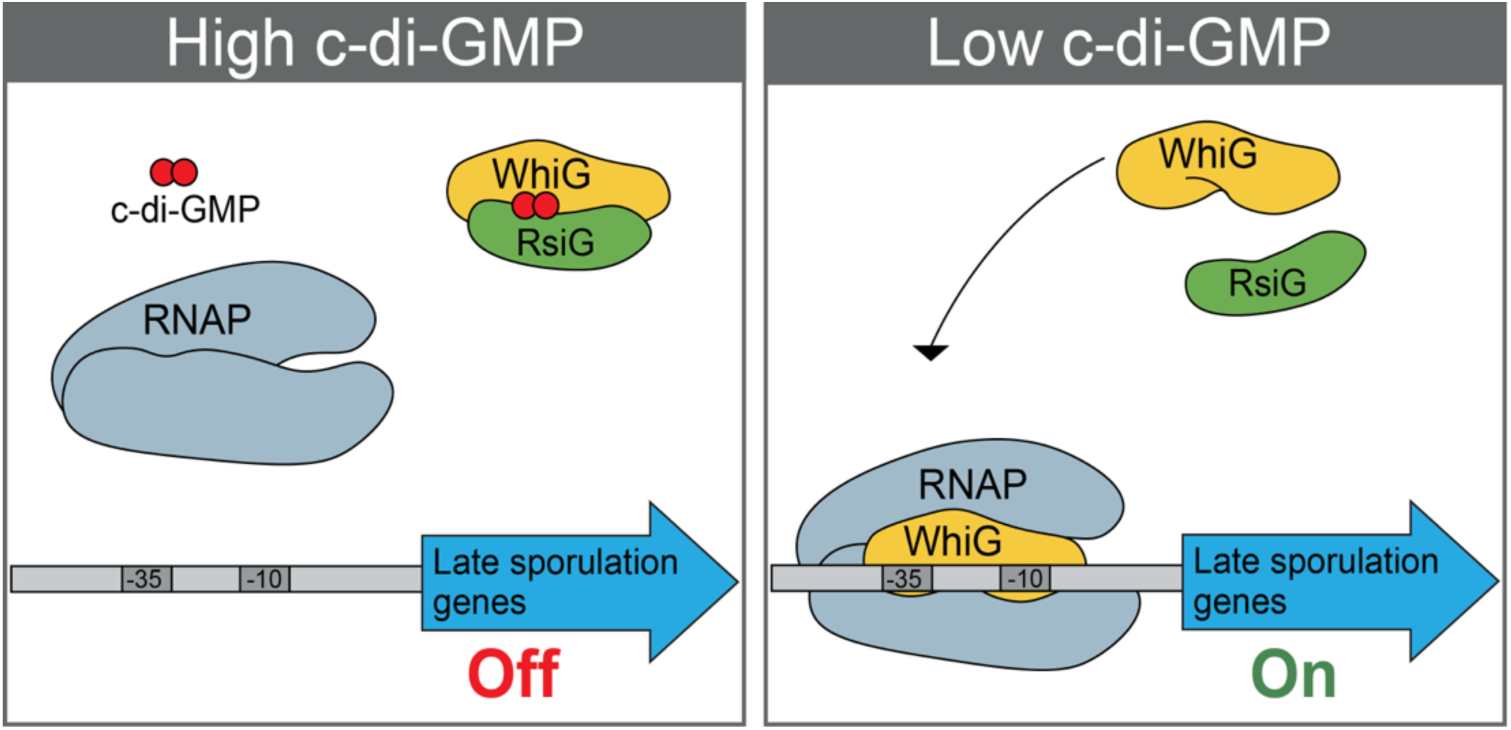
Schematic representation of c-di-GMP-dependent regulation of the RsiG-WhiG complex in *S. venezuelae.* (Left) At elevated intracellular c-di-GMP levels, anti-α RsiG binds WhiG via a c-di-GMP intercalated dimer, sequestering WhiG and preventing its association with RNA polymerase. (Right) At lower intracellular levels of c-di-GMP, the RsiG-(c-di-GMP)_2_-WhiG complex disassociates, freeing WhiG to associate with RNA polymerase and direct transcription of late sporulation genes.

RsiG is not homologous to any previously known c-di-GMP effector and its c-di-GMP-binding motif is unique. We recently showed that RsiG distribution is limited to Phylum Actinomycetota, where it is most conserved in the families Streptomycetaceae, Geodermatophilaceae, and Pseudonocardiaceae (17). E(X)_3_S(X)_2_R(X)_3_Q(X)_3_D signatures were found in a majority of RsiG sequences, suggesting that binding the second messenger is likely to be a conserved feature of these proteins. In the genus *Rubrobacter*, which descends from the most basal branch of the Actinomycetota (17), RsiG homologs possess only one copy of the c-di-GMP-binding motif. We showed that the single-motif RsiG from *Rubrobacter radiotolerans* must dimerize in order to form a complex with WhiG, and the crystal structure of the (RsiG)_2_-(c-di-GMP)_2_-WhiG complex from this species confirmed these homologs bind two molecules of c-di-GMP in a similar manner to *S. venezuelae* RsiG (17). These findings indicate that RsiG functions as a cognate anti-σ factor for WhiG in phylogenetically distant actinobacterial species.

To date, WhiG represents the only example of a σ factor known to be directly controlled by c-di-GMP (18). Phylogenetically, WhiG belongs to the σ^28^ family of alternative factors that regulate flagellum biosynthesis in diverse bacteria, including FliA in members of Phylum Pseudomonadota and SigD in members of Phylum Bacillota (19–21). Flagellar biosynthesis is subject to a regulatory hierarchy that ensures correct assembly of the complex appendage. During early stages of flagellar assembly, FliA/SigD is held inactive by a cognate anti-σ factor, FlgM (22–25). FlgM bears no homology to RsiG, is not known to bind c-di-GMP, and association with its cognate σ is regulated via a distinct mechanism. Specifically, following hook-basal body assembly, the hook length sensor FliK induces the secretion apparatus to change specificity (26) and FlgM is secreted (27–29), thereby releasing FliA/SigD. Once active, FliA/SigD is responsible for activating the expression of genes required for the late stages of flagellar biosynthesis, including the filament protein flagellin as well as genes involved in the chemosensory pathway (30, 31). Although most commonly associated with control of flagellar biosynthesis, FliA/SigD homologs have also been implicated in regulation of distinct functions including autolysis (32–34).

Within Phylum Actinomycetota, the function of WhiG homologs have been analyzed in two genera in addition to *Streptomyces*: *Rubrobacter* and *Actinoplanes*. In our previous analysis of *Rubrobacter* (RsiG)_2_-(c-di-GMP)_2_-WhiG, we showed that the WhiG homolog in this genus targets Type IV pilus biosynthesis, as well as several PDEs/DGCs, indicating the presence of regulatory feedback loops (17). Interestingly, the actinobacterial species *Actinoplanes missouriensis*, which feature a complex developmental life cycle that involve production of sporangia that release motile flagellated zoospores (35, 36), possess four paralogs of σ factors belonging to the WhiG/FliA family. An RNA-seq analysis by Hashigushi *et al*. (37) revealed that both developmental and motility-related genes are regulated by these σ factors. The sets of genes de-regulated by deletion of individual WhiG/FliA genes in *A. missouriensis* were only partially overlapping, indicating non-redundant functions for the paralogs. As a dedicated developmental σ factor, *S. venezuelae* WhiG has functionally diverged from the rest of the clade. Taken together, these observations suggest significant diversification of WhiG homolog function within the Actinomycetota.

So far, the role of WhiG homologs has been defined in a limited number of actinobacterial taxa and have not been subjected to systematic analysis. Because WhiG is known to be directly controlled by c-di-GMP in the phylogenetically distant actinobacterial genera *Streptomyces* and *Rubrobacter*, understanding the functional evolution of WhiG across the phylum will shed light on the processes controlled by c-di-GMP in these taxa. To examine the evolution of this σ factor, we searched a set of genomes representative of Phylum Actinomycetota to identify WhiG homologs. This analysis revealed that actinobacterial WhiG sequences can be classified into two distinct phylogenetic lineages, which we named WhiG1 and WhiG2, both of which are broadly distributed in diverse actinobacterial genera. WhiG1 sequences, which include those that are found in the zoospore-producing genus *Actinoplanes,* are co-conserved with the presence of the flagellar biosynthesis cluster, indicating that these homologs may regulate flagellar biosynthesis in diverse actinobacterial species. WhiG1 homologs are not able to form a complex with (c-di-GMP)-RsiG. By contrast, WhiG2 sequences, which include those found in *Streptomyces*, are likely to be regulated by c-di-GMP via RsiG partners. In support of these observations, our bioinformatic analysis predicts that chemotaxis and motility genes are frequently found in WhiG regulons. Along with these genes, we predict that WhiG homologs regulate genes involved in transcriptional regulation, c-di-GMP metabolism, and specialized metabolite biosynthesis. Overall, our analyses indicate a phylogenetically-predictable divergence in how actinobacterial WhiG homologs are regulated, as well as significant diversification in the function of WhiG through the evolution of Actinomycetota.

## Results

### Actinobacterial WhiG homologs form two distinct sub-clades

Our previous work assessing the distribution of the c-di-GMP-binding anti-σ factor RsiG, which harbors a unique c-di-GMP binding motif, revealed that it is found in diverse actinobacterial taxa and always co-occurs with a putative WhiG homolog (17). However, this previous study did not systematically address the distribution of WhiG homologs, and it has remained unclear if most actinobacterial WhiG homologs are likely to be regulated by RsiG partners. In addition, the identification of four WhiG paralogs in *A. missouriensis* (37) suggests that these proteins have been subject to expansions during evolution of the phylum, though the overall extent of this is unknown. Hence, to assess the distribution of WhiG within the Actinomycetota, we searched 673 reference/representative Actinomycetota genomes (Table S1) available at GenBank using the *S. venezuelae* WhiG (WhiG*_Sv_*) sequence as a query in a BLAST search. Identified sequences were then aligned with a set of reference sequences representing major σ^70^ families (38) (Table S2), and a phylogenetic tree was constructed. The smallest well-supported clade (>0.9) that included reference sequences from the FliA/SigD family was used to define the set of WhiG homologs (248 total). Of the 673 genomes analyzed, approximately 30% (187 genomes) possess a WhiG homolog. To refine the evolutionary relationships among the identified sequences, an actinobacterial WhiG phylogeny was then constructed (Fig 2). The initial bifurcation in the phylogeny reveals two distinct and well supported sub-clades, which were designated as the WhiG1 clade (96 sequences). and the WhiG2 clade (152 sequences, including WhiG*_Sv_*).

**Fig 2.**
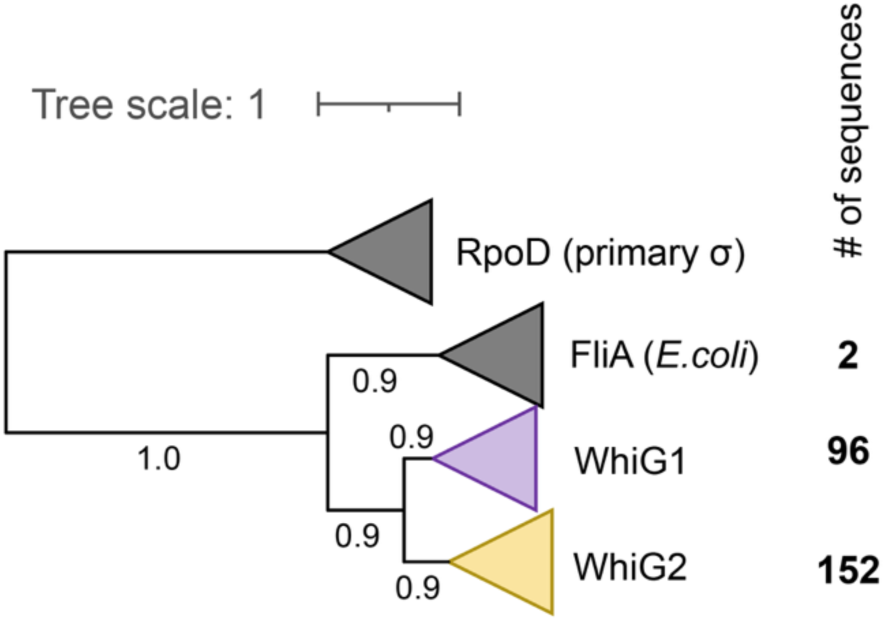
WhiG homologs cluster into two distinct and well supported subclades. Maximum likelihood phylogeny of WhiG homologs found in 673 reference Actinomycetota genomes. Bootstrap values are indicated at the respective nodes based on 100 replicates. The number of WhiG homologs found in each subclade is indicated to the right of each clade. Tree scale is substitutions per site.

### RsiG forms a complex with WhiG2, but not WhiG1 homologs

To assess the co-occurrence of the WhiG homologs with RsiG, we mapped the identified sequences onto a housekeeping phylogeny that reflects the evolutionary history of the 673 Actinobacterial genomes that were included in the search (17) (Fig 3). The presence of a WhiG2 homolog significantly correlated with the presence of a RsiG homolog (Mantel’s correlation coefficient *r* = 0.88, *p* = 0.001), but not with WhiG1 (Mantel’s correlation coefficient *r* = 0.09, *p* = 0.002) sequences. This co-distribution pattern suggested that WhiG2 homologs may be regulated by RsiG and c-di-GMP, as seen in the *S. venezuelae* model, whereas WhiG1 homologs may not form a complex with these molecules.

**Fig 3.**
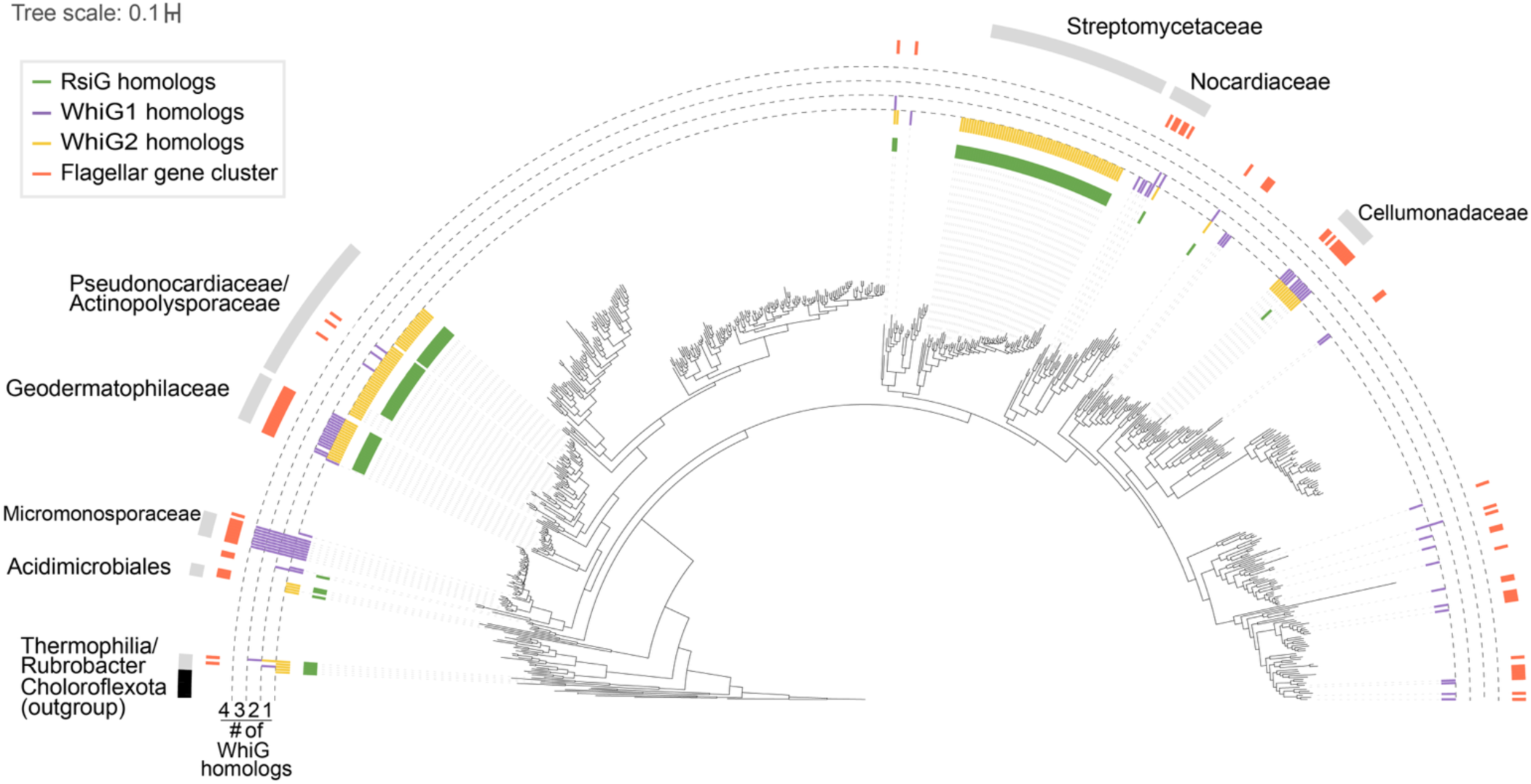
Distribution of WhiG1, WhiG2, RsiG homologs, and flagellar clusters across Phylum Actinomycetota. A maximum likelihood tree of 673 representative Actinomycetota genomes is shown, based on 37 concatenated housekeeping genes that were identified and aligned using PhyloSift (67). Sequences derived from five Chloroflexota genomes included as an outgroup (indicated in black). Genomes possessing an RsiG homolog are indicated by green bars. Presence of WhiG homologs is indicated by purple bars (WhiG1) and yellow bars (WhiG2); bar height reflects the number of WhiG homologs within each genome. Genomes possessing a flagellar gene cluster are indicated with orange bars. Taxonomic families with at least 50% of members containing a WhiG homolog are indicated by the gray arcs. Tree scale is substitutions per site.

To further explore the relationship between RsiG and the two clades of WhiG homologs, we employed Alphafold3 (AF3) modeling (39). AF3 predicts a high-confidence interaction (ipTM = 0.80) between the *S. venezuelae* RsiG (RsiG*_Sv_*) and WhiG*_Sv_* proteins (where high confidence is defined as an ipTM score ≥0.80). We therefore sought to predict these interactions in a representative genome from the family *Geodermatophilaceae*, where RsiG, WhiG1, and WhiG2 co-occur with high frequency (Fig 3). Members of this taxonomic family are frequently isolated from soil environments, have been described to grow as rods or rudimentary hyphae, and produce flagellated motile spores (40). An AF3 model of the RsiG and WhiG2 homologs encoded in the genome of *Geodermatophilus obscurus* DSM 43160 (RsiG*_Go_* and WhiG2*_Go_*), the type species of the genus (41), predict a high confidence interaction (ipTM 0.86), while the model of RsiG*_Go_* with the WhiG1 clade homolog from the same species (WhiG1*_Go_*) does not predict complex formation (ipTM 0.14) (Fig 4A,B). To validate these predictions *in vivo*, we assayed these proteins for interaction using a bacterial adenylate cyclase two-hybrid (BACTH) system. This confirmed that RsiG*_Go_* and WhiG2*_Go_* interact, in striking contrast to RsiG*_Go_* and WhiG1*_Go_*, which do not (Fig 4C).

**Fig 4.**
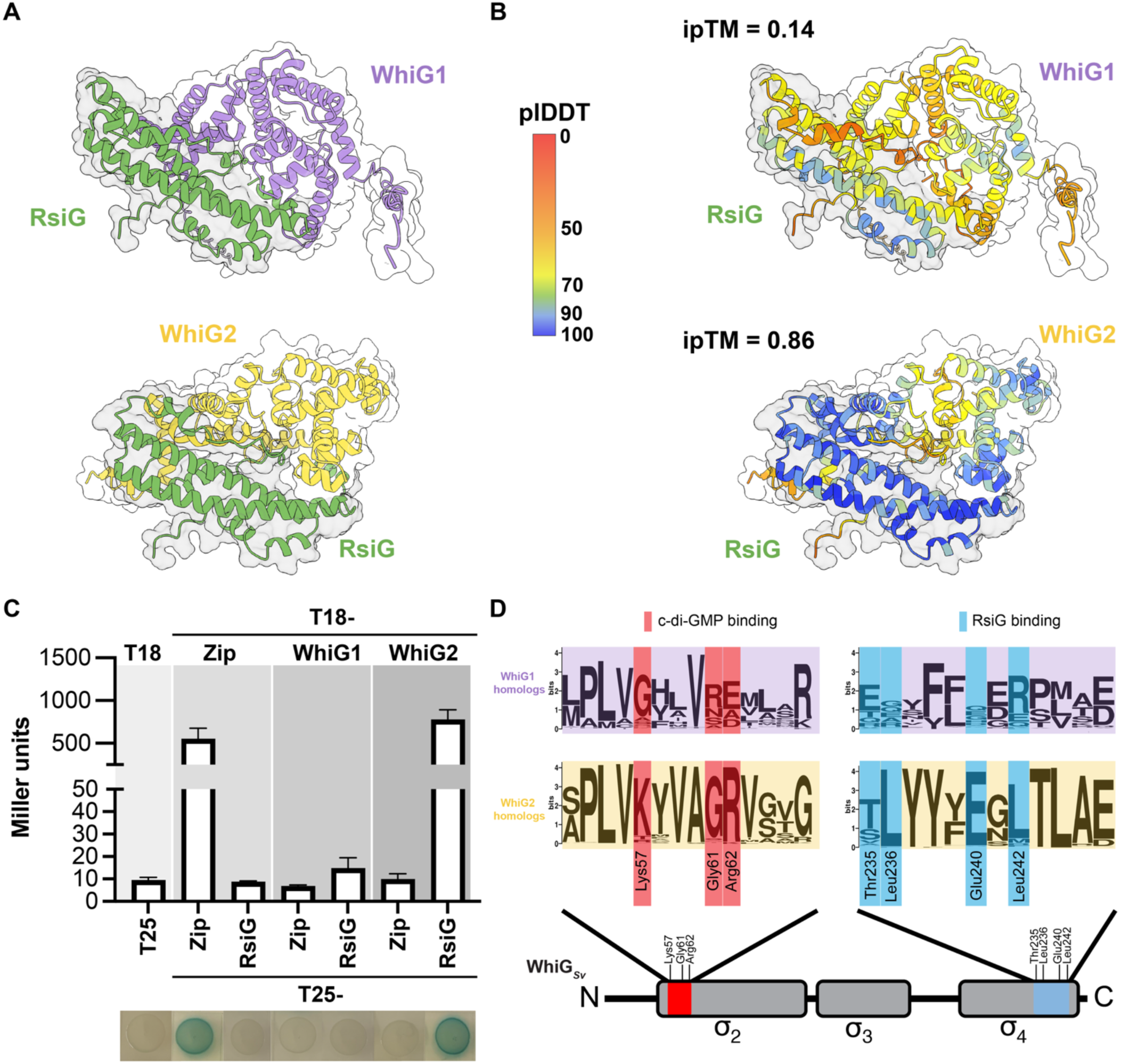
RsiG homologs interact only with WhiG2 homologs. (A) Alphafold3 models of RsiG*_Go_* (green) in complex with (top) WhiG1*_Go_* (purple) and (bottom) WhiG2*_Go_* (yellow). (B) Alphafold3 models of the interaction of RsiG*_Go_* with (top) WhiG1*_Go_* and (bottom) WhiG2_Go_ colored based on the structural confidence measure predicted lDDT (local distance difference test). Confidence in each complex is indicated by interface predicted template (ipTM) score. (C) Bacterial two-hybrid analysis to investigate the interaction between RsiG*_Go_* with WhiG1*_Go_* and WhiG2*_Go_*. The listed pairs of constructs were transformed into the BACTH reporter strain *E. coli* BTH101. Strains expressing fusions of both adenylate cyclase domains to the leucine zipper domain of GCN4 (Zip) served as a positive control, whereas strains expressing fusions of one adenylate cyclase domain to a Zip domain and the other to RsiG*_Go_*, WhiG1*_Go_,* or WhiG2*_Go_* served as negative controls. Results are the average of three replicate cultures derived from the same single colony. Error bars represent SEM. (D) Schematic depiction of WhiG domain organization below sequence logos depicting amino acid sequence conservation in the two clades of WhiG homologs at key sites within the α_2_ and α_4_ domains. Residues critical to the RsiG*_Sv_*-(c-di-GMP)_2_-WhiG*_Sv_* complex are noted below sequence logos. Residues that are critical for the interaction of WhiG*_Sv_* with c-di-GMP are highlighted in red and residues involved in the interaction of WhiG*_Sv_* with RsiG are highlighted in blue. Two separate multiple sequence alignments of WhiG1 homologs and WhiG2 homologs were used to generate the sequence logos using Weblogo (76).

To examine if our findings using the *G. obscurus* proteins are likely to extend to most members of the WhiG1 and WhiG2 clades, we assessed the conservation of residues critical for the interaction between the σ factor and the c-di-GMP ligand, as well as between the σ factor and anti-σ factor, by generating separate multiple sequence alignments for each clade. The structure of the *S. venezeulae* RsiG-(c-di-GMP)-WhiG complex revealed that three residues from the WhiG*_Sv_* α_2_ domain make direct contacts with the c-di-GMP dimer: Lys57, Gly61, and Arg62 (10). Conservation of Gly61, which packs against one of the ribose groups of c-di-GMP, is likely particularly important for the interaction, as any other side chain would cause a steric clash (10). Sequence logos made using these alignments illustrate that indeed, all three residues are well conserved among WhiG2, but not WhiG1 homologs (Fig 4D). Similarly, residues in the α_4_ domain of WhiG*_Sv_* (including Thr235, Leu236, Tyr239, Glu240, and Leu242), that make direct contacts with RsiG*_Sv_* (10) are well conserved in the alignment of the WhiG2 clade, but is highly variable in the WhiG1 clade (Fig 4D). Taken together, results of the co-distribution analysis, AF3-predicted structures, BACTH assay, and conservation of residues required for complex formation strongly suggest that RsiG, when present, is likely to physically interact with and regulate the activity of WhiG2 clade members via c-di-GMP but is unlikely to regulate WhiG1 clade homologs.

### WhiG1 sequences are co-distributed with the flagellar biosynthesis cluster

Notably, we almost always observe at least one WhiG1 homolog encoded in representative/reference genomes of the genera *Actinoplanes* and *Geodermatophilus* (Fig 3), both of which are known to have members that produce flagellated zoospores (35, 40). Based the phylogenetic relationship of WhiG with flagellar σ factors (i.e*. Salmonella* FliA and *Bacillus* SigD) (19) and recent findings that WhiG homologs in *A. missouriensis* control flagellar biosynthesis (37), we hypothesized that WhiG1 homologs may be involved in control of motility. To examine the co-occurrence of flagellar genes and WhiG1 in Actinomycetota, we conducted a cblaster (42) search of the set of actinobacterial representative genomes using the *A. missouriensis* 431 (NBRC 102363) cluster (43) as query (Table S3) and mapping the presence of the cluster onto the housekeeping phylogeny (Fig 3). We detected the presence of the flagellar cluster in 12% of genomes, with the highest conservation of the cluster observed in the genera *Actinoplanes* (Micromonosporaceae)*, Geodermatophilus* (Geodermatophilaceae), and *Cellulomonas* (Cellulomonoadaceae). This distribution of the flagellar gene cluster is largely in agreement with a previous analysis of the evolution of flagella in Actinomycetota (44). Strikingly, in 81% of cases where we observed a flagellar cluster, a WhiG1 clade homolog was also present (Mantel’s correlation coefficient *r* = 0.84, *p* = 0.001). This was not the case for WhiG2 clade homologs, which are present in only 46% of genomes with a flagellar cluster (Mantel’s correlation coefficient *r* = 0.14, *p* = 0.001). Furthermore, 62% of WhiG1 clade homologs are encoded within or near to the flagellar cluster itself (defined as being encoded within five genes of a gene within the cluster). These observations suggest that WhiG1 clade homologs may be functionally similar to the FliA/SigD family members found outside of Phylum Actinomycetota.

### Bioinformatic prediction of actinobacterial WhiG homolog regulons

To gain further insights into the functional diversification of these σ factors, we performed a bioinformatic search to identify likely WhiG target promoters. We relied on three previous studies of individual actinobacterial genomes to define a consensus sequence for this search. In the first of these, the WhiG2 target sequence in *S. venezuelae* was identified using ChIP-seq (10) and found to be highly similar to the well-established consensus (45) for the FliA/SigD family. Second, the previous analysis of *R. radiotolerans* used bioinformatic prediction combined with 5’ triphosphate end capture transcription start site mapping to predict 12 WhiG2 target sites (17). Finally, the WhiG1 target sequence in *A. missouriensis* was recently predicted by RNA-seq (37). From these analyses, we noticed that for both WhiG2 (reported consensus -35 T(A/C)AA; N16; -10 GC<u>CGA</u>TAA) and WhiG1 (reported consensus -35 CTCA; N16-17; -10 GC<u>CGA</u>ACT), the -35 region is highly conserved. In addition, the central CGA of the -10 (underlined) is also highly conserved. The most variability is seen in the -10 region excluding this central CGA. We therefore performed a bioinformatic search for these consensus sequences in the intergenic regions of genomes with a WhiG homolog. Positive hits were required to lie within 200 bp of a downstream start codon and allowed one mismatch, which could be in the -10, and no mismatches allowed in the -10 central CGA. Because a high degree of variability was observed for the final three nucleotides of the -10 among confirmed WhiG/FliA targets (10, 17, 37), we did not include these bases in our consensus queries. We limited this search to taxa within Actinomycetota with high conservation of WhiG homologs (defined as genera with at least two representatives in which a WhiG homolog is found in >80% of genomes). Given the high similarity between the two consensus sequences used as query, we next attempted to infer a WhiG1-specific regulon. Based on the strong correlation between WhiG1 and the flagellar cluster, we reasoned that this clade may specifically target flagellar genes. However, when we examined predicted binding motifs within flagellar clusters, the WhiG1 and WhiG2 query sequences were both present and overlapped extensively (Fig S1). Therefore, when both WhiG1 and WhiG2 are present in a single genome, we were unable to predict individual regulons for each homolog.

To analyze the identified targets, we constructed a sequence similarity network (46, 47), where protein sequences sharing ≥35% identity are connected via edges (Fig 5, Fig S2, Table S4). Conserved components of WhiG regulons are therefore grouped into clusters in the network. The resulting sequence similarity network is comprised of 131 total clusters. Notably, one of these, cluster 26, is comprised entirely of WhiG homologs, indicating that in some species, WhiG regulates its own promoter. Strikingly, the known *S. venezuelae* WhiG targets WhiI (Vnz_28820) and WhiH (Vnz_27205) were found in two of the largest clusters (clusters 5 and 6). Both of these clusters are primarily comprised of hits from Streptomycetaceae genomes (Fig 5, Fig S2), suggesting that these are highly conserved as WhiG targets within the family. Thirteen additional clusters are exclusively found in *Streptomyces* genomes, which may represent previously unknown and/or species-specific targets. Clusters comprised of genes found only in genomes from a single taxonomic family were also observed for Geodermatophilaceae, Micromonosporaceae, Pseudonocardiaceae, and Cellumonadaceae. In contrast, several clusters, including some of the largest clusters, are comprised of targets derived from genomes representing considerable taxonomic diversity. For example, the largest cluster in the network (cluster 1) has representatives from eight distinct actinobacterial families (Fig 5).

**Fig 5.**
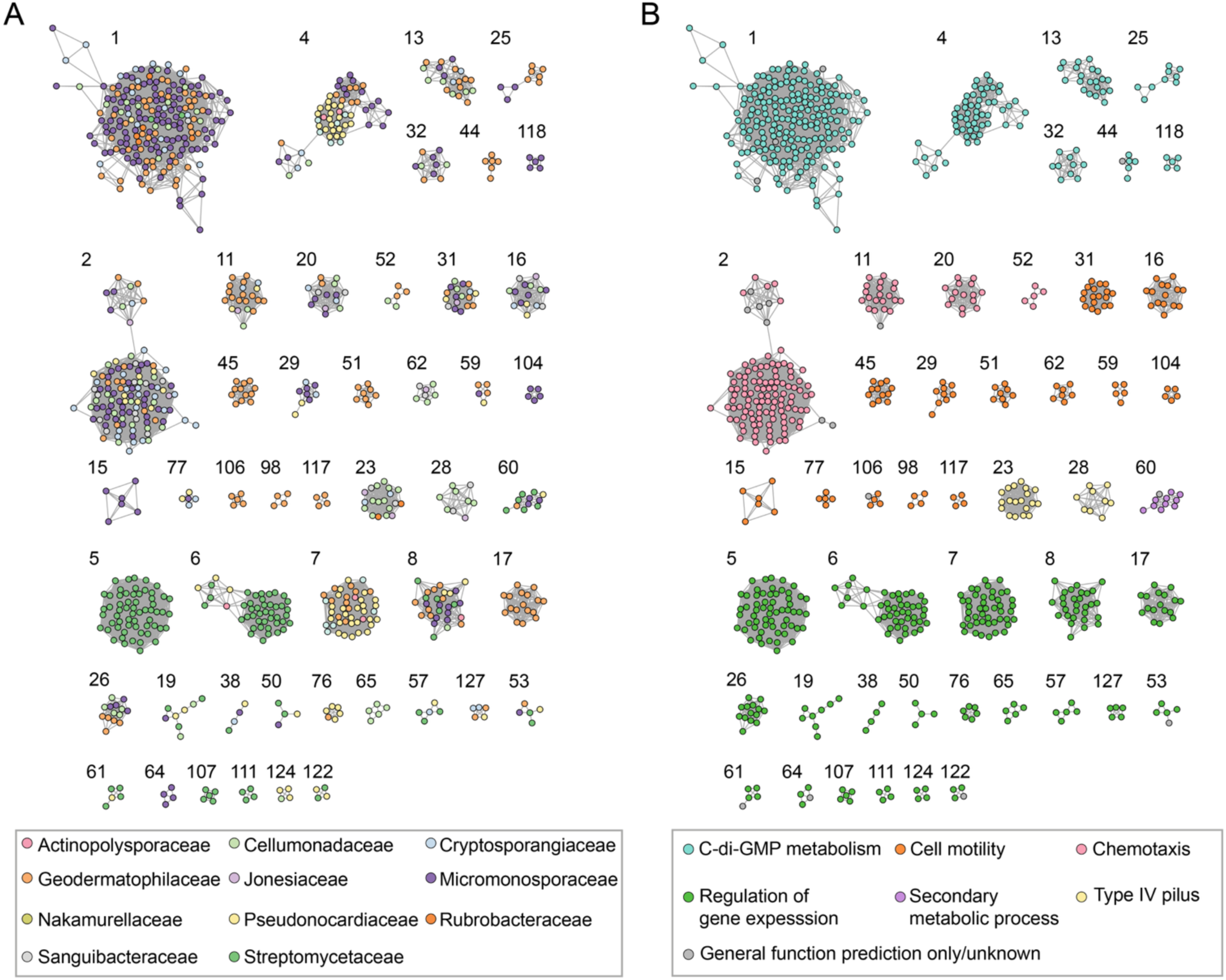
Subset of sequence similarity network of the predicted phylum-wide WhiG regulon. WhiG promoters (as defined in Methods) were identified in genera with at least two representatives in our dataset and in which WhiG homologs were present in >80% of genomes. Genes located immediately downstream of these promoters were subjected to all-versus-all BLAST and clustered using EFI-EST (46). The resulting network was filtered to include categories of biological interest (discussed in text) and visualized in Cytoscape version 3.10. 3 (82). A version of this network including all predicted target genes can be found in Fig S2. Clusters are numbered according to yFiles organic layout algorithm. Nodes represent individual genes, and edges denote BLAST pairwise similarity scores <1E-35. Nodes are colored according to taxonomic family (A) or associated biological process based on GO terms (B).

Functional predictions of each of the putative targets was performed using Gene Ontology (GO) assignments (Table S5), which revealed significant diversification of WhiG regulons. In addition, we generated HMM profiles from the sequences within each cluster. These HMM profiles were then searched against the *S. venezuelae* genome (Table S6) to identify homologs in a well-characterized reference system. Of clusters with a predicted biological process, most were involved in regulation of gene expression. Interestingly, using the HMM profile constructed using sequences from cluster 7, the top hit in *S. venezuelae* was BldM (Vnz_22005). BldM is an orphan atypical response regulator that in *Streptomyces* forms a heterodimer with WhiI, and together the two proteins control expression of approximately 22 genes (13). BldM is predicted to be a WhiG target in genomes representing disparate actinobacterial families, but not in any *Streptomyces* genomes. Outside of *Streptomyces*, it is unknown if BldM and WhiI form heterodimers. Additional targets that are predicted to be involved in regulation of gene expression include clusters where a majority of members are annotated as response regulators (cluster 5, which includes WhiI, cluster 6, which includes WhiH, and clusters 8, 17, 57, 64, 124), σ factors (cluster 26, which includes WhiG, and cluster 127), WhiB-like (cluster 107), LacI-family (clusters 19, 50), helix-turn-helix motif-containing (clusters 65, 61), as well as transcriptional regulators belonging to the TetR (cluster 111), AfsR (cluster 122), IclR (cluster 76), and ROK (cluster 53) families.

In addition to regulation of gene expression, one of the most conserved functional predictions we observed were the enzymes responsible for c-di-GMP metabolism. Seven clusters within our network (clusters 1, 4, 13, 25, 32, 44, 118) comprised of 307 putative target genes are predicted to function as DGC/PDE enzymes (Fig 5). These putative targets are found in eight distinct actinobacterial families, making c-di-GMP metabolism the most widespread function of WhiG targets across the Phylum. Despite this, we only observed two WhiG binding sites in front of genes encoding PDEs/DGCs in the family Streptomycetaceae, two EAL-domain containing proteins in the species *Streptomyces autolyticus* and *Streptomyces melanosporofaciens* (Fig 5, Table S4). In addition, for most genomes included in the analysis that possess genes for flagellar biosynthesis, we observed putative WhiG binding sites embedded within these gene clusters. Frequently, this occurs multiple times within a flagellar gene cluster, indicating that WhiG homologs potentially regulate the transcription of multiple operons in these clusters (Fig 6). Additional notable biological process categories identified include genes predicted to be involved in chemotaxis (four clusters) in members of seven distinct actinobacterial families (Fig 5, Table S4), and Type IV pilus biosynthesis pathways (two clusters) in members of the Cellumonadaceae, Cryptosporangiaceae, Sanguibacteraceae, Rubrobacteraceae, and Jonesiaceae (Fig 5). Genes involved in specialized metabolite biosynthesis (one cluster) are also predicted targets in several families, including the Micromonosporaceae, Pseudonocardiaceae, Geodermatophilaceae, and Streptomycetaceae. These targets include genes that are predicted to have core functions in specialized metabolite biosynthesis, including non-ribosomal peptide synthetases and polyketide synthases (Table S4).

**Fig 6.**
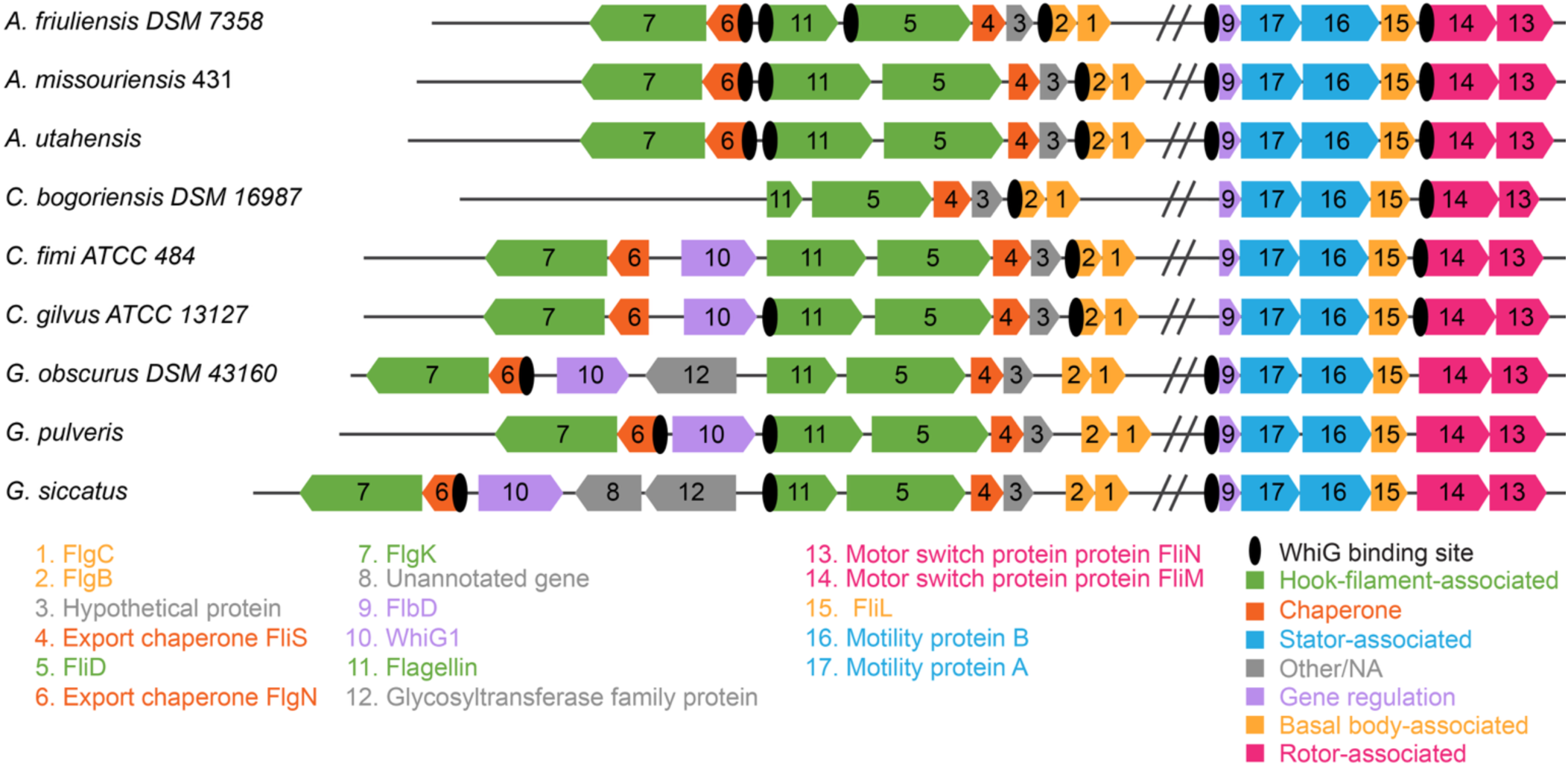
Predicted WhiG target promoters in flagellar gene cluster of *Actinoplanes friuliensis* DSM 7358, *Actinoplanes missouriensis* 431, *Actinoplanes utahensis*, *Cellulomonas bogoriensis* DSM 16987, *Cellulomonas fimi* ATCC 484, *Cellulomonas gilvus* ATCC 13127, *Geodermatophilus obscurus* DSM 43160, *Geodermatophilus pulveris*, and *Geodermatophilus siccatus*. Genes in the flagellar gene cluster are shown as schematics with the predicted gene products listed below with the same color coding. 24 matches to WhiG promoter sequence (defined in methods) were identified bioinformatically in the intergenic regions of aforementioned genomes.

## Discussion

Despite being broadly distributed in bacteria, to date the only known member of the flagellar-associated family of α factors known to be directly controlled via c-di-GMP are those found in Phylum Actinomycetota (17), exemplified by WhiG from the genus *Streptomyces*. The work presented here reveals that actinobacterial WhiG homologs can be classified into at least two distinct groups based on phylogenetic inference. WhiG1 homologs, are unable to associate with RsiG partners and have retained the ancestral association with the flagellar biosynthesis cluster. WhiG2 homologs, which include all of those found in the genus *Streptomyces*, are likely largely regulated by RsiG partners via c-di-GMP. We further demonstrate that the function of WhiG homologs are likely to be significantly more diverse than previously appreciated, showing how this regulator has been co-opted to control distinct functions in members of Phylum Actinomycetota.

The bifurcation of the actinobacterial WhiG phylogeny into two sister clades suggests that an ancient duplication event occurred during early evolution of the Actinomycetota. The co-distribution of most WhiG2 homologs with RsiG, which was not observed for WhiG1 homologs, suggested that regulation by RsiG partners may be specific to the WhiG2 clade. Indeed, in *G. obscurus*, a combination of AF3 modeling and bacterial two hybrid assays confirmed interaction between RsiG and WhiG2 is conserved in this species, but not with WhiG1. Key residues for the interaction with RsiG or c-di-GMP are highly conserved among WhiG2, but not WhiG1 homologs, which indicates that specific interaction of RsiG with WhiG2 is likely to extend beyond *G. obscurus* to most members of this clade. In contrast, WhiG1 homologs are unlikely to be regulated via RsiG partners, even when these two proteins co-occur in the same organisms. The identity of a putative anti-σ partner for WhiG1 homologs remains unclear, especially given the notable absence of FlgM, the cognate anti-σ of canonical flagellar α factors FliA/SigD, in almost all flagellar clusters found in Actinomycetota (44). The actinobacterial ancestor is predicted to have been motile and the flagellar cluster was likely largely inherited vertically throughout Phylum Actinomycetota, though loss events occurred along many lineages (44). Therefore, FlgM was likely lost as a part of the flagellar cluster during early evolution of the Phylum, and its absence may indicate an alternative, as-yet unidentified mechanism for regulation of the late stages of flagellar biosynthesis in these species.

The species that encode WhiG homologs comprise considerable morphological diversity and include filamentous and non-filamentous, motile and non-motile species. So far, the function of these homologs has been explored in a limited number of species that are relatively scattered throughout Phylum Actinomycetota. Our analysis of the role of WhiG therefore significantly expands on previously known roles for these σ factors. Owing to the similarity of published WhiG2 (10) and WhiG1 (37) consensus sequences, at this time it is unclear if these are responsible for the regulation of distinct or overlapping regulons when they co-occur in the same organism. However, our analysis has significantly expanded our understanding of the functional diversity of WhiG homologs in Phylum Actinomycetota. Regardless of the extent of regulon overlap, when WhiG1 and WhiG2 homologs co-occur they are likely to be regulated by distinct input signals, mediated by c-di-GMP_2_-RsiG (for WhiG2) and by an as-yet unidentified mechanism (for WhiG1).

We observed a spectrum of taxon-specificity for predicted WhiG target genes, with some targets found in genomes spanning the actinobacterial phylogeny, and other targets restricted to individual taxonomic families. One of the most abundant functional categories we identified were target genes predicted to encode DGCs/PDEs, the enzymes responsible for synthesis and degradation of c-di-GMP itself. These target genes were found in genomes that are very broadly phylogenetically distributed, representing all but one of the taxonomic families that were included in the search. Many of these predicted targets are from genomes where only a (c-di-GMP)-RsiG-regulated WhiG2 homolog is found, suggesting the presence of regulatory feedback loops responsible for control of c-di-GMP metabolism. A notable exception to the broad phylogenetic distribution of DGC/PDE targets are the streptomycetes: only two of these DGCs/PDEs are from in *Streptomyces* genomes, which represent just 3% of the total genomes belonging to Streptomycetaceae found in our search set. Interestingly, another transcription factor that is directly regulated by c-di-GMP, called BldD, is known repress transcription of multiple DGCs in *Streptomyces* species (16, 48–50). BldD is a repressor that must form a complex with c-di-GMP to bind its target genes (48, 51). In *S. venezuelae,* BldD is active during vegetative growth, while WhiG2 becomes active during the maturation of aerial hyphae into spores. This suggests that for *Streptomyces,* there may be more direct regulation of c-di-GMP via expression of genes encoding DGCs during earlier stages of the life cycle.

Genes involved in motility and chemotaxis were frequently identified as predicted WhiG targets. This includes most species included in our search that possess a flagellar gene cluster. The conservation of components of chemosensory pathways and structural proteins of the flagellar filament in these regulons suggests that the role of actinobacterial WhiG homologs is likely to be required for later stages of flagellar biosynthesis, similar to the role of SigD/FliA in the Bacillota and Pseudomonadota (30, 31). From studies of these phylogenetically distant phyla, c-di-GMP itself is known to directly regulate the process of motility at multiple layers including regulation of gene expression, flagellar assembly, motor rotation, and chemotaxis (52). For example, in *Pseudomonas aeruginosa,* c-di-GMP directly controls expression of flagellar genes via the transcription factor FleQ. c-di-GMP binding reduces ATPase activity of FleQ, resulting in a switch from activation of flagellar genes to activation of biofilm matrix genes (53–56). In some bacterial species, c-di-GMP is also known to control flagellar assembly by binding to and inhibiting the ATPase activity of a subunit of the export apparatus, FliI (57). In *E. coli*, c-di-GMP can also directly control the flagellar motor output itself via the PilZ-domain containing YcgR. When in complex with c-di-GMP, YcgR acts as a brake on cell motility (58, 59). In addition to control of motor output, in some species c-di-GMP directly regulates chemotaxis through MapZ, which represses the methyltransferase activity of CheR1, thereby altering the response to stimuli (60). This interaction requires c-di-GMP binding via the PilZ domain of MapZ. Thus, the direct and multi-layered regulation of motility via c-di-GMP appears to be broadly conserved across diverse bacterial phyla.

Our analysis also revealed the presence of WhiG binding sites within clusters responsible for the biosynthesis of Type IV pili. Type IV pili are surface appendages that are involved in diverse functions including competence, twitching motility, surface sensing, and microcolony formation (61, 62). Notably, in *A. missouriensis*, Type IV pili are present on zoospores in addition to flagella, and are required for adhesion to hydrophobic solid surfaces (63). Predicted binding sites within these clusters were previously confirmed in *A. missouriensis* (37) and *R. radiotolerans* (17). Our analysis suggests that regulation of Type IV pilus biosynthesis by WhiG is conserved across the genera *Actinoplanes* and *Rubrobacter*. In addition, we predict that Type IV pilus biosynthesis is under the control of WhiG for members of Jonesiaceae, Sanguibacteraceae, Cellulomonadaceae, and Cryptosporangiaceae. Like motility, Type IV pilus biosynthesis in Gram-negative bacteria is known to be a c-di-GMP-regulated process, and c-di-GMP can affect multiple levels of this regulation. For example, c-di-GMP inhibits transcription of the major pilin gene, *pilA,* in *Myxococcus* (64). c-di-GMP is also known to interact directly with pilus machinery in *Vibrio cholera* via interaction with the extension ATPase MshE (65).

One of the most frequent functional categories of genes that we predict to be a part of WhiG regulons are transcription factors. In several cases, this includes WhiG itself (cluster 26), which is consistent with observations in *Salmonella* (66) and *Bacillus* (33) that FliA/SigD control the transcription of *fliA/sigD*. Additionally, our analysis revealed that WhiI is a target of WhiG2 homologs in almost all (97%) of the *Streptomyces* genomes included in our search. WhiH homologs were also found as predicted targets in a majority (67%) of *Streptomyces* genomes. In *S. venezuelae,* WhiH is known to control a large regulon of late sporulation genes, including the genes responsible for biosynthesis of the volatile compounds geosmin and 2-methylisoborneal, which have been shown to be important for spore dispersal (67). Similarly, *S. venezuelae* WhiI in complex with BldM controls a regulon of ∼40 sporulation genes (13). Thus, by activating transcription of *whiH* and *whiI, Streptomyces* WhiG homologs are likely to indirectly control large sets of genes involved in late stages of sporulation. Interestingly, BldM homologs also appear in our SSN (cluster 7), although none of these hits were found in *Streptomyces* genomes. The separation of BldM and WhiI expression has a clear function in *Streptomyces*: prior to WhiI expression, BldM forms a homodimer and regulates genes that are required for earlier stages of the life cycle. Once *whiI* expression is activated by WhiG, BldM forms a heterodimer and controls a second set of genes required for spore maturation (13). Outside of *Streptomyces*, it is unknown if BldM and WhiI are able to form a functional heterodimer. The appearance of BldM as a major target outside of *Streptomyces* provides an example of rewiring of related transcriptional networks through the evolution of Actinomycetota. Additional uncharacterized transcription factors that we have identified during our analysis may represent novel downstream regulators in the WhiG-regulated pathways, such as development and motility, and will be the subject of future study.

Given the high degree of conservation of the signature c-di-GMP-binding motif(s) among RsiG homologs (17), it is likely that most of these homologs are able to bind the second messenger. Thus, the identification of WhiG2 regulons are, by extension, predictions of the processes that are directly regulated by c-di-GMP signaling. Interestingly, there does appear to be some functional overlap between WhiG1 and WhiG2, as we frequently found predicted WhiG binding sites in front of PDEs and DGCs in genomes that possess single homologs belonging to either clade. Overall, this work points to significant diversification of WhiG regulation and function over the course of actinobacterial evolution. Further work is needed to understand whether WhiG1 and WhiG2 regulons overlap when the two proteins co-exist.

## Materials and Methods

### Actinomycetota phylogeny

A search set of 673 Actinomycetota genomes were chosen based on annotation as ‘reference’ or ‘representative’ at GenBank and use in a previous analysis of RsiG distribution (17), along with ten genomes from Phylum Chloroflexota that serve as an outgroup (Table S1). The Chloroflexota genome *Anaerolinea thermophila* UNI-1 was used to root the tree. Amino acid sequences of 37 conserved housekeeping genes were automatically identified, aligned, and concatenated using Phylosift (68). Model selection was performed using SMS (69) implemented at http://www.atgc-montpellier.fr/phyml/ (70) which resulted in selection of a LG substitution model with γ-distributed rate variation between sites. Phylogenetic reconstruction was performed by RAxML version 8.2.10 (71) with 100 rapid bootstraps replicates to assess node support. The tree was visualized and formatted using iTOL (72). Taxonomic assignments were based on the taxonomy database maintained by the NCBI (https://www.ncbi.nlm.nih.gov/Taxonomy/Browser/wwwtax.cgi).

### WhiG homolog identification and alignment

WhiG homologs were identified by a BLAST search (e-value cutoff = 0.001) of the same set of 673 Actinomycetota annotated genomes using the WhiG*_Sv_* sequence as a query. This resulted in a set of 3,836 initial hits that were aligned using MAFFT (73) with a set of 16 reference sequences that includes representatives from all σ^70^ groups (38) (Table S2). 1,769 sequences that did not align well and were not reciprocal hits to WhiG*_Sv_* in a BLAST search of the *S. venezuelae* genome were removed from the analysis. The resulting multiple sequence alignment was then used to construct an approximately-maximum-likelihood phylogeny using FastTree 2 (74). The smallest well-supported clade (>0.9) that included all reference sequences from the σ^28^ family was used to define the set of WhiG homologs (248 total). This set of phylogeny-confirmed actinobacterial WhiG homolog sequences were then re-aligned using CLUSTAL Omega (75). Model selection was performed using ModelTest-NG (76) which resulted in selection of a LG substitution model with γ-distributed rate variation between sites. Phylogenetic reconstruction was performed by RAxML version 8.2.10 (71) with 100 rapid bootstraps replicates to assess node support. The tree was visualized and formatted using iTOL (72). To visualize conservation of residues involved in interaction with c-di-GMP or RsiG, WhiG1 (n =96) or WhiG2 (n =152) homologs were extracted from the alignment and sequence logos generated using Weblogo (77).

### Flagellar Cluster Identification

To identify genomes that possessed the genetic potential to produce a flagellum, a search for the flagellar gene cluster in our local database of 673 reference Actinomycetota genomes was performed using cblaster (42) with default parameters (minimum query coverage = 50%, minimum sequence identity = 30%, and a maximum distance between any two hits = 20 kbp). 33 flagellar genes (Table S3) from *Actinoplanes missouriensis* 431 (43) were used in the query. Genomes were considered to possess a flagellar gene cluster if they contained at least 30% of the query genes (e-value cutoff = 0.01).

### Statistical analysis

Partial Mantel tests with the Pearson method were used to determine correlations between the presence of WhiG1 or WhiG2 homologs and RsiG homologs or the flagellar cluster while controlling for the influence of phylogenetic distance. Mantel tests were performed in R version 4.5.1 (78) using the package vegan (79) and set to 999 permutations. A phylogenetic distance matrix was obtained from the maximum likelihood tree of 673 representative actinobacterial species using the “cophenetic.phylo” function of the package ape (80).

### Protein modeling

Protein complex predictions of WhiG1*_Go_* and WhiG2*_Go_* with RsiG generated using AlphaFold3 (39). Confidence in the complexes were measured by interface predicted template modeling (ipTM) score (39). ChimeraX version 1.8 (81) was used to generate structural representations.

### Bacterial strains, plasmids, and media

Strains, plasmids, and oligonucleotides used in this study are listed in Table S7. *E. coli* strain DH5α was used for plasmid propagation and grown on LB or LB agar at 37 °C. Where required for selection, the following antibiotics were added to growth media: 100 μg/mL carbenicillin and/or 50 μg/mL kanamycin.

### Bacterial adenylate cyclase two-hybrid assays

*E. coli* codon-optimized versions of the *rsiG*, *whiG1*, and *whiG2* genes from *G. obscurus* DSM 43160 were generated by gene synthesis (Twist Biosciences). These were PCR-amplified with primers JD07-JD18 (Table S7) and assembled into pUT18 and pKT25 (10, 17, 82) digested with XbaI and KpnI to create plasmids pJD001-pJD012 (Table S7). *E. coli* BTH101 was then co-transformed with the “T18” and “T25” fusion plasmids. pUT18 and pKT25 empty constructs were used as a negative control. β-galactosidase activity was assayed in biological triplicate.

### Prediction of WhiG regulons

In genomes possessing a WhiG homolog, we searched intergenic regions within 200 bp upstream of a start codon for established WhiG promoter consensus sequences. Genomes with only WhiG2 homologs were searched with T(A/C)AA-N_16_-GCCGA (10, 17) as query, those with only WhiG1 homologs were searched with CTCA-N_16-17_-GCCGA (37) as query, and those containing both homologs were searched with both motif queries. The -35 region was restricted to exact matches to T(A/C)AA or CTCA. The -10 region was required to contain CGA at positions 3-5 and up to one mismatch was allowed in positions 1-2. This search resulted in 7,199 matches. The amino acid sequences of the downstream genes of the matched motifs were uploaded to EFI-EST (47) to generate a sequence similarity network (SSN) (edge e-value cutoff = 1.0E-5). The generated SSN was further filtered to retain clusters comprised of >3 nodes, resulting in 131 clusters encompassing 1490 nodes (Fig S2, Table S4) Sequence similarity networks were visualized using Cytoscape (83). Clusters were numbered according to yFiles organic layout algorithm.

Functional annotation of WhiG targets was performed by Blast2GO (84), which employs the Gene Ontology Consortium (85, 86) database to assign GO terms based on existing annotation associations. GO terms were categorized into broad functional categories using ancestor charts and parental terms (Table S5). Nodes with broad or generic annotations (e.g., GO:0016787 hydrolase activity) were analyzed with InterProScan (87) to identify functional domains and then sorted into a functional annotation category (e.g., EAL domain sorted to “c-di-GMP metabolism”). Nodes still lacking a specific annotation or with no annotation available were classified as “General function prediction only/unknown.” In addition, predicted WhiG target sequences were binned by cluster number and aligned using MUSCLE (88). Resulting alignments were used to create hidden Markov model (HMM) profiles with “hmmerbuild” from the HMMER suite (hmmer.org, version 3.4). These profiles were queried against the *Streptomyces venezuelae* genome (GCA_001886595.1) using “hmmsearch” of the HMMER suite.

To illustrate the conservation of WhiG targets in the flagellar gene cluster, 9 representative species from three genera with high levels of WhiG1 conservation were selected. FlaGs (Flanking Genes) (89) was used to identify sets of homologous flanking genes and create a graphical visualization of the flagellar gene cluster. Genomes were aligned according the two most frequently targeted flagellar genes, flagellin (n =18) and FlbD (n = 18).

## Author contributions

J.D.D., Y.V.B., and K.A.G. designed research; J.D.D., G.C., J.J.C., and K.A.G performed research and analyzed data, J.D.D. and K.A.G. wrote the manuscript.

## Acknowledgements

We thank Tory Hendry, Neil Holmes, Tiffany Zarella, John Helmann, Daniel Buckley, and Mark Buttner for helpful discussions and comments on the manuscript. This work was funded by Canada 150 Research Chair in Bacterial Cell Biology (to Y.V.B.) and NIH R35GM156858 (to K.A.G.).

