## Supplementary Material for "Regulatory divergence and functional diversification of a c-di-GMP-controlled sigma factor in Actinomycetota"

**Table S1.** Reference Actinomycetota and Chloroflexota genomes
See separate file

**Table S2.** Reference $\sigma^{70}$ sequences used in phylogenetic analysis

| **UniProt Accession** | **Protein** | **Organism** |
|---|---|---|
| F2R2N1 | SigR | *Streptomyces venezuelae* |
| P0AEM6 | FliA | *Escherichia coli* |
| P10726 | SigD | *Bacillus subtilis* |
| Q9KQD4 | FliA | *Vibrio cholera* |
| P0AGB3 | RpoH | *Escherichia coli* |
| P06574 | SigB | *Bacillus subtilis* |
| P07860 | SigF | *Bacillus subtilis* |
| P12254 | SigK | *Bacillus subtilis* |
| P13445 | RpoS | *Escherichia coli* |
| H8F0N6 | SigE | *Mycobacterium tuberculosis* |
| P45684 | RpoS | *Pseudomonas aeruginosa* |
| P00579 | RpoD | *Escherichia coli* |
| P52324 | RpoD | *Caulobacter vibrioides* |
| P17531 | RpoD | *Myxococcus xanthus* |
| Q06909 | CarQ | *Myxococcus xanthus* |
| Q06198 | AlgU | *Pseudomonas aeruginosa* |

**Table S3.** Flagellar genes in *Actinoplanes missouriensis* 431 used as queries for cblaster search in Actinomycetota genomes

| **Gene** | **Gene Product** |
|---|---|
| AMIS76150 | FliW |
| AMIS76160 | FlgL |
| AMIS76170 | FlgK |
| AMIS76180 | FlgN |
| AMIS76190 | FliC |
| AMIS76200 | FliD |
| AMIS76210 | FliS |
| AMIS76220 | Unknown |
| AMIS76230 | FlgB |
| AMIS76240 | FlgC |
| AMIS76250 | FliE |
| AMIS76260 | FliF |
| AMIS76270 | FliG |
| AMIS76280 | FliH |
| AMIS76290 | FliI |
| AMIS76300 | Unknown |
| AMIS76310 | LytA |
| AMIS76320 | FliK |
| AMIS76330 | Unknown |
| AMIS76340 | Unknown |
| AMIS76350 | FlhA |
| AMIS76360 | FlhB |
| AMIS76370 | FliR |
| AMIS76380 | FliP |
| AMIS76390 | FliO |
| AMIS76400 | FliN |
| AMIS76410 | FliM |
| AMIS76420 | FliL |
| AMIS76430 | MotB |
| AMIS76440 | MotA |
| AMIS76450 | FlbD |
| AMIS76460 | FlgE |
| AMIS76470 | FlgD |

**Table S4.** Predicted WhiG targets in Actinomycetota
See separate file

**Table S5.** GO term assignment sorted into categories of biological associated process

| GO term | Biological associated process |
|---|---|
| GO:0052621 (diguanylate cyclase activity) | **c-di-GMP metabolism** |
| GO:0071111 (cyclic-guanylate-specific phosphodiesterase activity) | |
| GO:0003924 (GTPase activity) | catalytic activity (**General function prediction only/unknown**) |
| GO:0016491 (oxidoreductase activity) | |
| GO:0016740 (transferase activity) | |
| GO:0016787 (hydrolase activity) | |
| GO:0016853 (isomerase activity) | |
| GO:0020037 (heme binding) | |
| GO:0140096 (catalytic activity, acting on a protein) | |
| GO:0140097 (catalytic activity, acting on DNA) | |
| GO:0140098 (catalytic activity, acting on RNA) | |
| GO:0048870 (cell motility) | **Cell motility** |
| GO:0005737 (cytoplasm) | cellular component (**General function prediction only/unknown**) |
| GO:0005886 (plasma membrane) | |
| GO:0016020 (membrane) | |
| GO:0044283 (small molecule biosynthetic process) | |
| GO:0110165 (cellular anatomical structure) | |
| GO:0009987 (cellular process) | |
| GO:1901135 (carbohydrate derivative metabolic process) | |
| GO:0006575 (modified amino acid metabolic process) | |
| GO:0006935 (chemotaxis) | **Chemotaxis** |
| GO:0003677 (DNA binding) | molecular function (**General function prediction only/unknown**) |
| GO:0005515 (protein binding) | |
| GO:0035438 (cyclic-di-GMP binding) | |
| GO:0140657 (ATP-dependent activity) | |
| NA | NA (**General function prediction only/unknown**) |
| GO:0005975 (carbohydrate metabolic process) | **Primary metabolic process** |
| GO:0006520 (amino acid metabolic process) | |
| GO:0006629 (lipid metabolic process) | |
| GO:0030163 (protein catabolic process) | |
| GO:0036211 (protein modification process) | |
| GO:0006355 (regulation of DNA-templated transcription) | **Regulation of gene expression** |
| GO:0140110 (transcription regulator activity) | |

| | |
|---|---|
| GO:0044550 (secondary metabolite biosynthetic process) | **Secondary metabolite biosynthesis** |
| GO:0006412 (translation) | **Translation** |
| GO:0045182 (translation regulator activity) | |
| GO:0006810 (transport) | **Transport** |
| GO:0043683 (type IV pilus assembly) | **Type IV pilus** |

**Table S6.** HMM profiles of WhiG regulon clusters used to searched *Streptomyces venezuelae* genome
See separate file

**Table S7.** Strains, plasmids, and primers used in this study

| Strains | Relevant genotype/notes | Reference |
|---|---|---|
| DH5α | F– φ80*lac*ZΔM15 Δ(*lac*ZYA-*argF*)U169 *rec*A1 *end*A1 *hsd*R17(rK–,mK+) *pho*A *sup*E44 λ– *thi*-1 *gyr*A96 *rel*A1 | Invitrogen |
| BTH101 | F– *cya*-99 *ara*D139 *gal*E15 *gal*K16 *rps*L1 (*Strr*) *hsd*R2 *mcr*A1 *mcr*B1 | Karimova et al., 1998 |

| Plasmid | Relevant genotype/comments | Source |
|---|---|---|
| pKT25 | Two-hybrid plasmid, N-terminal cyaAT25 fusion ($Kan^R$) | Karimova et al., 1998 |
| pKNT25 | Two-hybrid plasmid, C-terminal cyaAT25 fusion ($Kan^R$) | Karimova et al., 1998 |
| pUT18 | Two-hybrid plasmid, C-terminal cyaAT25 fusion ($Amp^R$) | Karimova et al., 1998 |
| pUTI8C | Two-hybrid plasmid, N-terminal cyaAT25 fusion ($Amp^R$) | Karimova et al., 1998 |
| pKT25-zip | A derivative of pKT25 in which the leucine zipper of GCN4 is genetically fused in frame to the T25 fragment | Karimova et al., 1998 |
| pUT18C-zip | A derivative of pUT18C in which the leucine zipper of GCN4 is genetically fused in frame to the T18 fragment | Karimova et al., 1998 |
| pJD001 | pUT18 carrying $whiG2_{Go}$ | This work |
| pJD002 | pUT18C carrying $whiG1_{Go}$ | This work |
| pJD003 | pUT18 carrying $whiG1_{Go}$ | This work |
| pJD004 | pUT18C carrying $whiG1_{Go}$ | This work |
| pJD005 | pUT18 carrying $rsiG_{Go}$ | This work |
| pJD006 | pUT18C carrying $rsiG_{Go}$ | This work |
| pJD007 | pKT25 carrying $whiG2_{Go}$ | This work |
| pJD008 | pKNT25 carrying $whiG2_{Go}$ | This work |
| pJD009 | pKT25 carrying $whiG1_{Go}$ | This work |
| pJD010 | pKNT25 carrying $whiG1_{Go}$ | This work |
| pJD011 | pT25 carrying $rsiG_{Go}$ | This work |
| pJD012 | pNT25 carrying $rsiG_{Go}$ | This work |

| Primers | 5’ sequence |
|---|---|
| JD07 | CTGCAGGTCGACTATGACCAAAGCGGCGAT |
| JD08 | TGAATTCGAGCTCGGAGCGATATCAGCCAGTTTG |
| JD09 | CTGCAGGTCGACTATGCGTTCTAGCGCGAGC |
| JD10 | TGAATTCGAGCTCGGCGCGCTACGACCGGTAGC |
| JD11 | CTGCAGGTCGACTATGACCCCGGAACCGACC |
| JD12 | TGAATTCGAGCTCGGACGACCTTCATCAGCCAG |
| JD13 | GCTGCAGGGTCGGACTACCAAAGCGGCGATC |
| JD14 | TTCTTAGTTACTTAGGAGCGATATCAGCCAGTTTG |
| JD15 | GCTCGAGGGTCGACTATGCGTTCTAGCGCGAGC |
| JD16 | TTCTTAGTTACTTAGGCGCGCTACGACCGGTAGC |
| JD17 | GCTCGAGGGTCGACTATGACCCCGGAACCGACC |
| JD18 | TTCTTAGTTACTTAGGACCTTCATCAGCCAG |

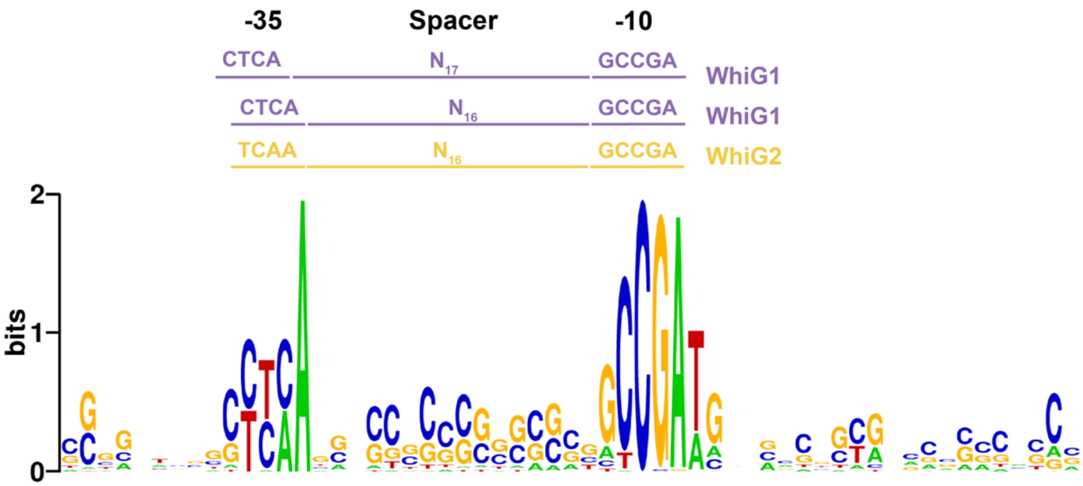


**Fig S1** Sequence logo of predicted WhiG binding motifs in flagellar gene clusters. Predicted WhiG binding motifs sequences found in the flagellar gene cluster of genomes possessing both WhiG1 and WhiG2 homologs were used to create a multiple sequence alignment, which was then used to generate the sequence logo using Weblogo (1). Overlapping WhiG1 and WhiG2 promoter queries are depicted above the sequence logos.

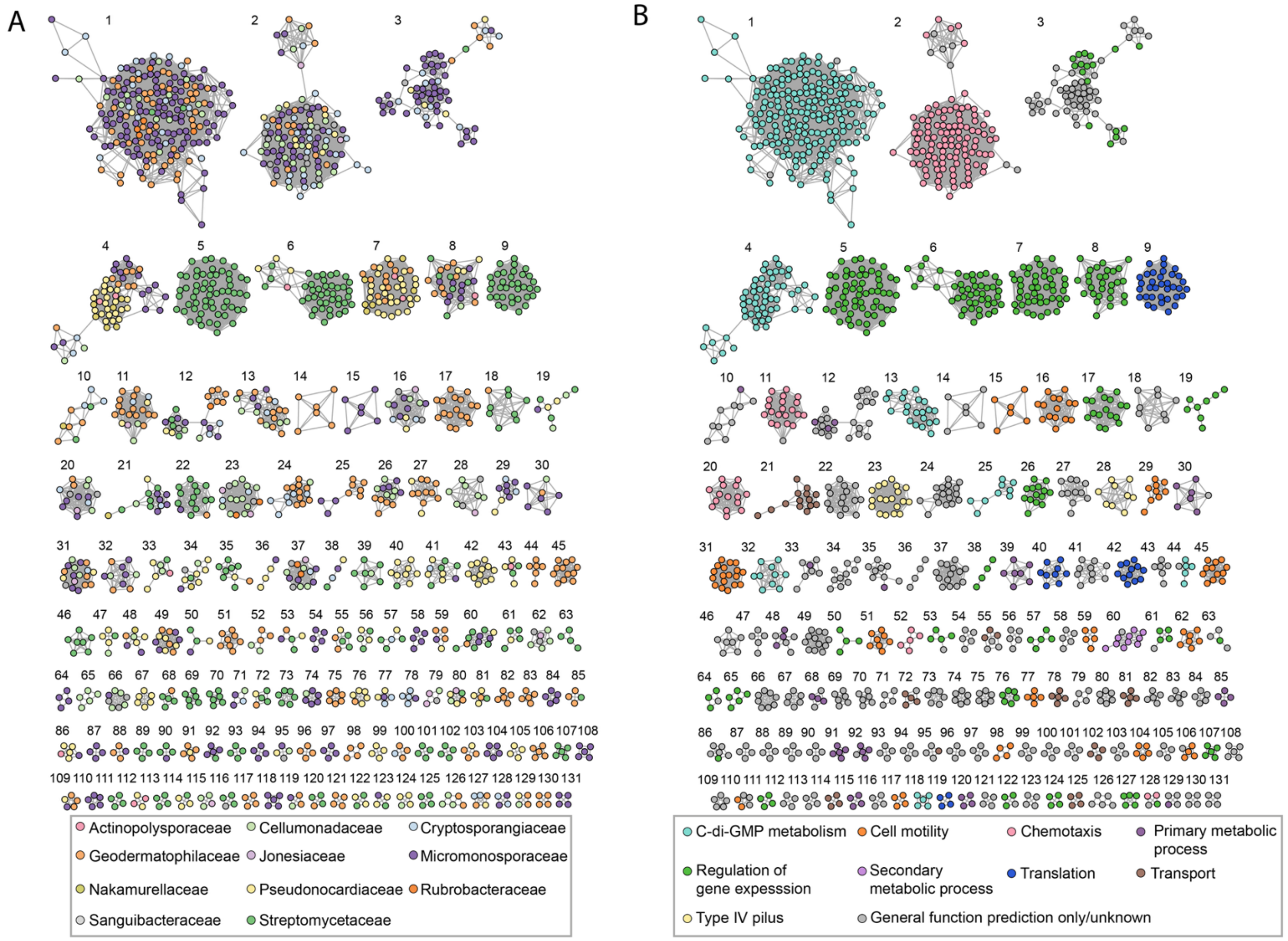


**Fig S2** Full sequence similarity network of the predicted phylum-wide WhiG regulon. WhiG promoters were identified in genera with at least two representatives in our dataset and in which WhiG homologs were present in >80% of genomes. Genes located immediately downstream of these promoters were subjected to all-versus-all BLAST and clustered using EFI-EST (2). The resulting network is visualized in Cytoscape version 3. 10.3 (3). Clusters are numbered according to yFiles organic layout algorithm (4). Nodes represent individual genes, and edges denote BLAST pairwise similarity scores <1E-35. Nodes are colored according to taxonomic family (A) or associated biological process based on GO terms (B).
